# Intermediate filaments EXC-2 and IFA-4 Maintain Luminal Structure of the Tubular Excretory Canals in *Caenorhabditis elegans*

**DOI:** 10.1101/277111

**Authors:** Hikmat I. Al-Hashimi, David H. Hall, Brian D. Ackley, Erik A. Lundquist, Matthew Buechner

## Abstract

The excretory canals of *Caenorhabditis elegans* are a model for understanding the maintenance of apical morphology in narrow single-celled tubes. Light and electron microscopy shows that mutants in *exc-2* start to form canals normally, but these swell to develop large fluid-filled cysts that lack a complete terminal web at the apical surface, and accumulate filamentous material in the canal lumen. Here, whole-genome sequencing and gene rescue show that *exc-2* encodes intermediate filament protein IFC-2. EXC-2/IFC-2 protein, fluorescently tagged via CRISPR/Cas9, is located at the apical surface of the canals independently of other intermediate filament proteins. EXC-2 is also located in several other tissues, though the tagged isoforms are not seen in the larger intestinal tube. Tagged EXC-2 binds via pulldown to intermediate filament protein IFA-4, which is also shown to line the canal apical surface. Overexpression of either protein results in narrow but shortened canals. These results are consistent with a model whereby three intermediate filaments in the canals, EXC-2, IFA-4, and IFB-1, restrain swelling of narrow tubules in concert with actin filaments that guide the extension and direction of tubule outgrowth, while allowing the tube to bend as the animal moves.

**Article Summary:** The *C. elegans* excretory canals form a useful model for understanding formation of narrow tubes. *exc-2* mutants start to form normal canals that then swell into fluid-filled cysts. We show that *exc-2* encodes a large intermediate filament (IF) protein previously not thought to be located in the canals. EXC-2 is located at the apical (luminal) membrane, binds to another IF protein, and appears to be one of three IF proteins that form a flexible meshwork to maintain the thin canal diameter. This work provides a genetically useful model for understanding the interactions of IF proteins with other cytoskeletal elements to regulate tube size and growth.

## INTRODUCTION

Polarized cells form tubular structures ubiquitously in living organisms (LUBARSKY and KRASNOW 2003; IRUELA-ARISPE and DAVIS 2009; MARUYAMA and ANDREW 2012). Tubes vary in width, length, and in mechanism of formation (LUBARSKY and KRASNOW 2003; SIGURBJORNSDOTTIR *et al.* 2014). The mechanism by which a narrow biological tube grows and maintains a uniform diameter throughout the lifespan of an organism is poorly understood, however. The excretory system of the nematode *Caenorhabditis elegans* provides a useful model of “seamless” (no intracellular adherence junctions) single-celled tubular structures (SUNDARAM and BUECHNER 2016) such as vertebrate capillaries or the tip cells of the *Drosophila* trachea (LUBARSKY and KRASNOW 2003). The core excretory system consists of a large excretory cell plus a duct and pore cell (NELSON *et al.* 1983). The excretory cell, located beneath the pharynx, extends four hollow canals throughout the length of the worm roughly in the shape of the letter “H” (Fig. 1A, B). The canals collect and excrete excess water from the body through the duct and pore to regulate organismal osmolarity (NELSON and RIDDLE 1984). Worms with defects in excretory canal function exhibit pale bloated canals and bodies, and have less tolerance to high-salt environments (BUECHNER *et al.* 1999; WANG and CHAMBERLIN 2002; HAHN-WINDGASSEN and VAN GILST 2009).

**Figure 1.**
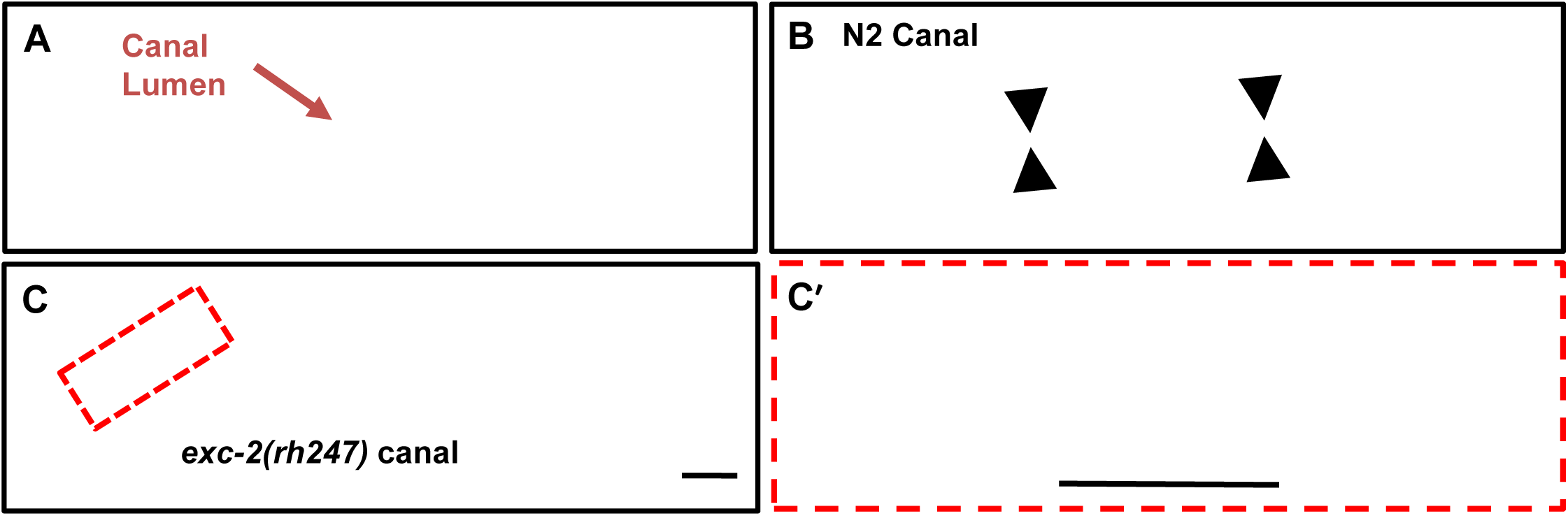
*exc-2* mutant shows short and cystic canals. (A) Diagram showing the excretory canals in wild-type *C. elegans* with two tubes (red apical surface, black basolateral surface, lumen in white) extended over the entire length of the worm and connected at the canal cell body. Canals extend from cell body in both directions anteriorward and posteriorward. (B) DIC image of excretory canal of wild-type worm (N2); arrows indicate narrow canal lumen with uniform diameter. (C) DIC image of *exc-2*(*rh247*) mutant shows canal extending to only ∼30% of the wild-type length. Area outlined in red is magnified in C’ to show the fluid-filled cysts accumulated throughout entire canal. Scale bars, 50µm.

In genetic screens, mutations in nine “*exc*” genes were discovered to affect canal structure to allow fluid-filled cysts to accumulate during canal extension during late embryogenesis and early first larval stage (BUECHNER *et al.* 1999). Other studies found similar mutations that affect additional tubular structures in the nematode, including the seamed single-cell excretory duct cell (JONES and BAILLIE 1995; MANCUSO *et al.* 2012; GILL *et al.* 2016; PU *et al.* 2017) and the multicellular intestine (BOSSINGER *et al.* 2004; ZHANG *et al.* 2011; CARBERRY *et al.* 2012; KHAN *et al.* 2013; ZHU *et al.* 2015; GEISLER *et al.* 2016). In the canal cell, proteins implementing tubule structure comprise apical cytoskeletal elements (MCKEOWN *et al.* 1998; PRAITIS *et al.* 2005; KHAN *et al.* 2013; KOLOTUEV *et al.* 2013; SHAYE and GREENWALD 2015), vesicular trafficking and exocyst proteins (TONG and BUECHNER 2008; MATTINGLY and BUECHNER 2011; ARMENTI *et al.* 2014; LANT *et al.* 2015; GRUSSENDORF *et al.* 2016), and ion and lipid transporters (BERRY *et al.* 2003; KHAN *et al.* 2013), among others.

Cytoskeletal components play an essential role in maintaining canal structure (SUNDARAM and COHEN 2017). Actin filaments are aligned over the apical surface of the canal lumen and docked to apical membrane via the ezrin/radixin/moesin homologue ERM-1 (GÖBEL *et al.* 2004) and the apical β_H_spectrin (PRAITIS *et al.* 2005), while mutations in the formin gene *exc-6* compromise nucleation of microtubules along the length of the canal (SHAYE and GREENWALD 2015). *C. elegans* contains eleven cytosolic intermediate filament (IF) proteins, plus one nuclear lamin protein (DODEMONT *et al.* 1994; KARABINOS *et al.* 2001). Three intermediate filaments proteins are highly expressed in the canal cell: IFC-2, IFA-4, and IFB-1 (SPENCER *et al.* 2011). Knockdown of the *ifb-1* gene causes cystic defects in both the canal and the multicellular intestine (WOO *et al.* 2004; KOLOTUEV *et al.* 2013).

While most of the original *exc* genes have been cloned (GRUSSENDORF *et al.* 2016; SUNDARAM and BUECHNER 2016), mutations in the *exc-2* gene cause particularly severe canal defects. In these mutants, the canal length is shortened by over half, the animals accumulate multiple cysts in the canals, and are sensitive to growth at low osmolarity (BUECHNER *et al.* 1999). Four alleles of this gene were discovered in the original screen, which suggested that it encodes a large protein. Here we report that *exc-2* encodes the intermediate filament IFC-2, and additionally found that mutations in the *ifa-4* intermediate filament gene also cause cystic canal defects similar to those of *exc-2* mutants. Overexpression of either *exc-2* or *ifa-4* results in shortened canals with small or no cysts. EXC-2 and IFA-4 proteins bind to each other and are located at the apical membrane of the canals. The position of EXC-2 at the apical membrane occurs independently of IFB-1 and IFA-4 function in the canals. These results indicate the importance of these three intermediate filaments in forming and maintaining the uniform diameter of the canals in this single-celled model of long, narrow tubular structure.

## MATERIALS AND METHODS

### DNA constructs and dsRNA synthesis

This study utilized two canal markers: pCV01 (used at 15ng/µl) contains the *gfp* gene driven by the canal-specific *vha-1* promoter; and pBK162 (used at 25ng/µl) contains the *mCherry* gene driven by the *exc-9* promoter. Fosmid WRM0630A_E08 was provided by the Max Planck Institute, Dresden, Germany. Genomic DNA was used for *exc-2* rescue and was prepared via PCR with LongRange enzyme (Qiagen, Venlo, NL) to amplify the full-length *ifc-2* gene, including 2kb upstream and 500bp downstream. The translational construct of *ifa-4* was made by ligating *ifa-4* cDNA in-frame with the *gfp* gene in plasmid pCV01.

Double-stranded RNA (dsRNA) constructs were synthesized via PCR-amplification of 250-350bp regions of selected exons in genes of interest (Supp. Table 1). A MEGAscript T7 kit (ThermoFisher, Waltham, MA) was used for transcription. For *ifb-1*, RNAi constructs were created by placing two constructs, each containing a complementary sequence corresponding to exon 4, under control of the canal-specific *vha-1* promoter.

**Table 1.**
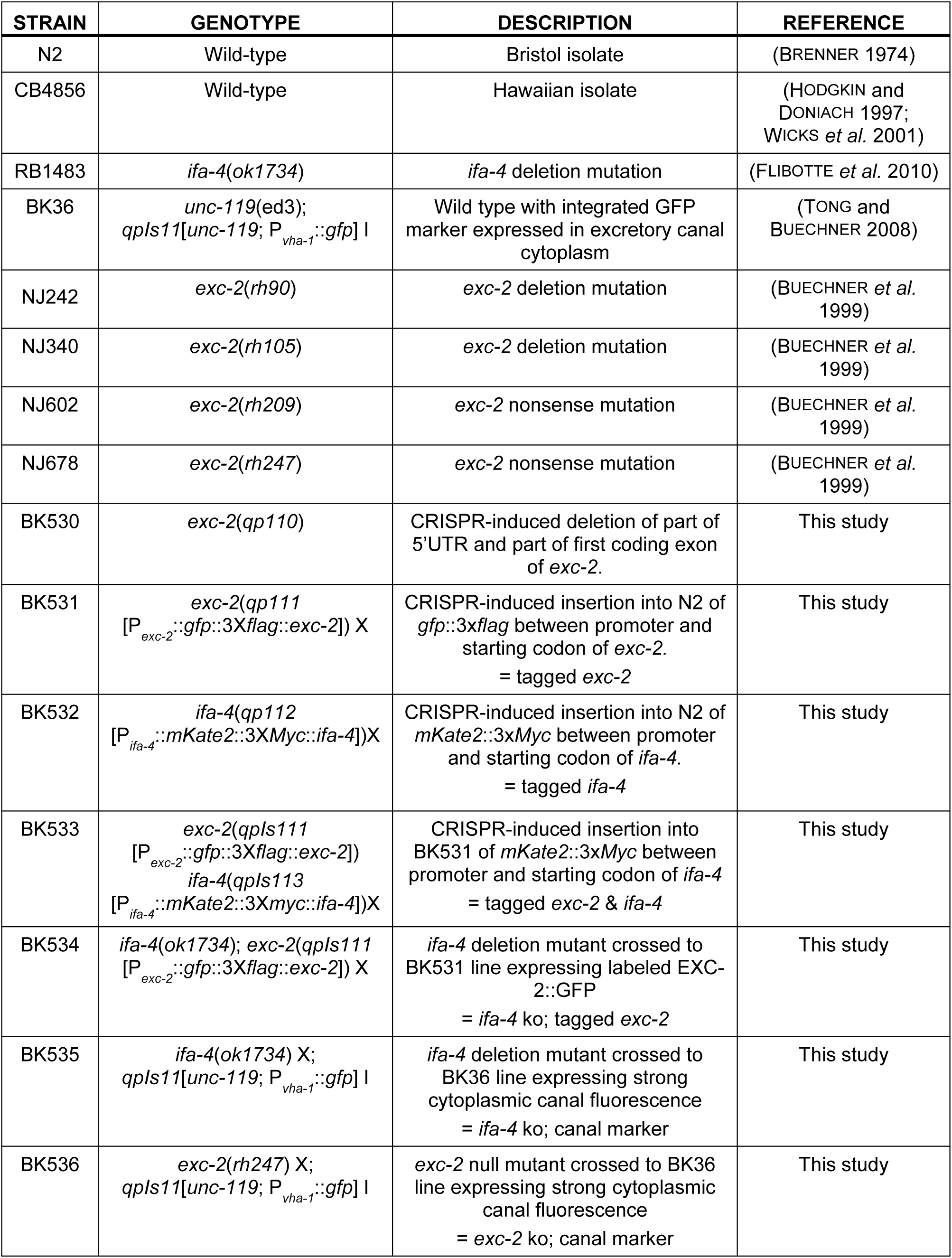
List of strains used in this study, with genotype descriptions.

| STRAIN | GENOTYPE | DESCRIPTION | REFERENCE |
| --- | --- | --- | --- |
| N2 | Wild-type | Bristol isolate | (BRENNER 1974) |
| CB4856 | Wild-type | Hawaiian isolate | (HODGKIN and DONIACH 1997; WICKS *et al.* 2001) |
| RB1483 | *ifa-4(ok1734)* | *ifa-4* deletion mutation | (FLIBOTTE *et al.* 2010) |
| BK36 | *unc-119(ed3); qpls11[unc-119; P<sub>vha-1</sub>::gfp]* I | Wild type with integrated GFP marker expressed in excretory canal cytoplasm | (TONG and BUECHNER 2008) |
| NJ242 | *exc-2(rh90)* | *exc-2* deletion mutation | (BUECHNER *et al.* 1999) |
| NJ340 | *exc-2(rh105)* | *exc-2* deletion mutation | (BUECHNER *et al.* 1999) |
| NJ602 | *exc-2(rh209)* | *exc-2* nonsense mutation | (BUECHNER *et al.* 1999) |
| NJ678 | *exc-2(rh247)* | *exc-2* nonsense mutation | (BUECHNER *et al.* 1999) |
| BK530 | *exc-2(qp110)* | CRISPR-induced deletion of part of 5'UTR and part of first coding exon of *exc-2*. | This study |
| BK531 | *exc-2(qp111 [P<sub>exc-2</sub>::gfp::3Xflag::exc-2])* X | CRISPR-induced insertion into N2 of *gfp::3xflag* between promoter and starting codon of *exc-2*.<br>= tagged *exc-2* | This study |
| BK532 | *ifa-4(qp112 [P<sub>ifa-4</sub>::mKate2::3XMyc::ifa-4])* X | CRISPR-induced insertion into N2 of *mKate2::3xMyc* between promoter and starting codon of *ifa-4*.<br>= tagged *ifa-4* | This study |
| BK533 | *exc-2(qpls111 [P<sub>exc-2</sub>::gfp::3Xflag::exc-2])*<br>*ifa-4(qpls113 [P<sub>ifa-4</sub>::mKate2::3Xmyc::ifa-4])* X | CRISPR-induced insertion into BK531 of *mKate2::3xMyc* between promoter and starting codon of *ifa-4*<br>= tagged *exc-2* & *ifa-4* | This study |
| BK534 | *ifa-4(ok1734); exc-2(qpls111 [P<sub>exc-2</sub>::gfp::3Xflag::exc-2])* X | *ifa-4* deletion mutant crossed to BK531 line expressing labeled EXC-2::GFP<br>= *ifa-4* ko; tagged *exc-2* | This study |
| BK535 | *ifa-4(ok1734)* X;<br>*qpls11[unc-119; P<sub>vha-1</sub>::gfp]* I | *ifa-4* deletion mutant crossed to BK36 line expressing strong cytoplasmic canal fluorescence<br>= *ifa-4* ko; canal marker | This study |
| BK536 | *exc-2(rh247)* X;<br>*qpls11[unc-119; P<sub>vha-1</sub>::gfp]* I | *exc-2* null mutant crossed to BK36 line expressing strong cytoplasmic canal fluorescence<br>= *exc-2* ko; canal marker | This study |

Constructs for CRISPR/Cas9-mediated mutation were created according to the method of the Goldstein laboratory (DICKINSON *et al.* 2015). The sgRNA constructs were made by amplifying plasmid pDD162 using primers containing a 20-base sequence specific for the gene of interest. The PCR product was self-ligated after treatment with T4 kinase. Goldstein group construct pDD282 was used for tagging *exc-2*, and pDD287 to tag *ifa-4*. Repair constructs were prepared through Gibson assembly of constructs containing the gene-specific tags in four overlapping amplified fragments, and ligated via NEBuilder® HiFi DNA Assembly Cloning Kit (NewEngland Biolabs, Ipswich, MA).

### Nematode genetics and genetic mapping

Strains of *C. elegans* are shown in Table 1. They were maintained on lawns of bacterial strain BK16 (a streptomycin-resistant OP50 strain) on NGM agar plates as described (SULSTON and HODGKIN 1988).

By means of complementation tests and deficiency mapping, *exc-2* was previously mapped to the left end of the X chromosome (BUECHNER *et al.* 1999). Strains of *exc-2* to be sequenced were each outcrossed to a wild-type Hawaiian isolate (CB4856) as described (MINEVICH *et al.* 2012). For each of four mutant allele strains, twelve F2 progeny homozygous for the *exc-2* mutation were selected, and grown to populations that were combined for whole-genome sequencing. Sequencing was completed at the Genome Sequencing Core at the University of Kansas. Genome data analysis was carried out to identify mutations in the expected genetic area by use of the Galaxy cloud-map website (https://usegalaxy.org).

Genetic rescue assays of *exc-2*(*rh247*) mutants were performed through co-injection into the gonad of carrier DNA pCV01, plus either Fosmid WRM0630A_E08, or PCR-amplified genomic *exc-2* DNA. Injected animals were allowed to lay eggs, which were screened for expression of the GFP-expressing carrier DNA. F1 progeny expressing GFP were examined for canal morphology.

Rescue of the *ifa-4* deletion mutant strain RB1483 was performed through gonad co-injection of an *ifa-4* cDNA construct at 40ng/µl together with marker plasmid pBK162. Injection of these *exc-2* and *ifa-4* constructs was also used to cause overexpression of the genes in wild-type worms. RNAi-knockdown of specific *exc-2* isoforms was accomplished through co-injection of forward and reverse RNA together with carrier pCV01 into gonads of young adult wild-type worms.

Complementation tests were carried out by mating male *exc-2*(*rh247*) to hermaphrodites of the BK530 (CRISPR’d *ifc-2* deletion) strain. Fluorescent hermaphrodite cross-progeny all showed the strong cystic canal phenotype of *exc-2* (n=30).

CRISPR/Cas9-mediated knock-in strains were created through injection of repair constructs from modifications to pDD282 and pDD287 CRISPR reagents (AddGene.org, Cambridge, MA) to make plasmids pBK301 and pBK302 in order to insert a fluorescent marker and epitope tag between the 5’UTR and coding region of *exc-2* and *ifa-4*, respectively. Selection of strains containing the constructs was performed on NGM plates containing 250 μg/ml hygromycin (Sigma-Aldrich, St. Louis). Heat-shock at 35C for 4-5 hours was used to activate removal of the selection cassette via self-excision. As the *exc-2* and *ifa-4* genes are located close together on the X chromosome, the doubly-tagged *exc-2*; *ifa-4* strain BK533 was created via CRISPR/Cas9-mediated tagging of BK531 (tagged *exc-2*) with pBK302.

### Microscopy and Canal Measurement

Worms were examined through a Zeiss Axioskop microscope with Nomarski (DIC) optics and epifluorescence. Animals were placed on 3% agarose pads in water and immobilized either through addition of Polybeads® polystyrene beads (Polysciences, Warrington, PA) or of sodium azide (35 mM). Non-confocal images were taken using an Optronics MagnaFire Camera. Some images of larger worms required 2 or 3 photographs that were “stitched” together to provide picture of the entire animal. Contrast on DIC images was uniformly enhanced over the entire image to increase clarity. For protein subcellular location, worms were examined using an Olympus Fluoview FV1000 laser-scanning confocal microscope. Lasers were set to 488nm excitation and 520nm emission (GFP), or 543nm excitation and 572nm emission (mKate2). All images were captured via FluoView optics (Olympus, Tokyo, JP) and collocation was analyzed using ImageJ software by drawing a straight line perpendicular to the length of the canal. Fluorescence plot profiles were then recorded and analyzed as shown in each image.

Electron microscopy was performed as described (BUECHNER *et al.* 1999). Young adult worms were cut and fixed in buffered glutaraldehyde and OsO_4_, encased in agar, dehydrated, and embedded in resin. Serial sections are ∼70 nm thick, and stained in uranyl acetate and lead citrate.

Canals were measured for length and cysts size as described (TONG and BUECHNER 2008). The length of each canal was scored between 0 and 4, where 4 indicates a full-length canal; 3 for canals that extended between the vulva and full-length; 2 for canals extending to the vulva; 1 for short canals that extended between the cell body and the vulva; 0 for canals that did not extend past the cell body. Cyst size was scored by measuring cyst diameter relative to worm body width. Large cysts have a diameter similar to that of the worm body width, medium cysts have a diameter of approximately half-worm width, and small cysts are smaller than half the worm body width. A 3x2 Fisher’s Exact Test was used to compare number of animals with short, medium-length, and full-length canals; and separately to compare number of animals with cysts, no cysts but canals with enlarged diameter, and normal-diameter canals.

### Biochemistry and Binding Assays

Worms were collected after growth on strain BK16 cultured on twelve 100mm plates of NGM medium until the bacterial lawn was consumed, then washed in M9 solution and frozen in liquid nitrogen until needed. Worm lysates were prepared through vortexing 500µl thawed worms in a mixture of 300µl dry 425-600 µm-diameter glass beads (Sigma-Aldrich, St. Louis, MO) and 700µl lysis buffer (0.5% NP40, 150mM NaCl, 50mM Tris pH 8.0, 0.5mM EDTA, 5% glycerol, 1mM DTT, and Protease-inhibitor tablet (Thermo Fisher Scientific, Waltham, MA)). The mixture was vortexed for 15min at maximum speed at 4^°^C, followed by 15 minutes on ice to cool down the samples, and again for another 15 minutes at 4^°^C. The lysate was then centrifuged in an Eppendorf centrifuge at maximum speed for 5 minutes at 4^°^C, and protein concentration of the supernatant was measured.

Protein samples for co-immunoprecipitation were incubated with pre-blocked (using 5% BSA) anti-FLAG® M2 magnetic beads (Sigma-Aldrich) at 4^°^C for 30 minutes. Samples were then washed thoroughly with PBS-T and eluted using 3X FLAG peptides (Sigma-Aldrich).

Samples for western blots were loaded onto Mini-PROTEAN® TGX^TM^ gels with a 4-20% gradient of bis-acrylamide (Bio-Rad, Hercules, CA). For immunoprecipitation samples, an equal volume was loaded in each well, while for lysate samples an equal amount of protein was loaded each well. Nitrocellulose membranes were blocked in 5% instant milk in TBS-T.

Tagged EXC-2 protein was detected via monoclonal anti-FLAG® M2 antibody (Sigma-Aldrich) at a final concentration of 1µg/ml in blocking buffer, and tagged IFA-4 protein was detected via HRP-bound anti-c-Myc antibody (Sigma-Aldrich) at a final concentration of 0.5µg/ml in blocking buffer.

### Data Availability

Sequences of *exc-2* mutant genes will be deposited at GenBank and to Wormbase.org. Strains of mutant alleles, and CRISPR/Cas9-modified strains of fluorescently and antigen-labeled *exc-2* and *ifa-4* will be made available through the *Caenorhabditis* Genetics Center, U. Minnesota (cgc.umn.edu). CRISPR Repair construct plasmids pBK301 and pBK302 are available upon request.

## RESULTS

Mutants in all alleles of *exc-2* show severely cystic canal phenotypes, with multiple fluid-filled cysts evident along the entire length of the greatly shortened canals (Fig. 1C and C’, Supp. Fig. 1). Canal length is only 30-40% of that of wild-type canals, and cysts have an average diameter 4 times as wide as that of a wild-type canal lumen. Electron microscopy images of *exc-2* canals show areas where the electron-dense actin-rich terminal web of the apical membrane has been thinned or lost (Fig. 2B and B’) in comparison with that of wild-type canal (Fig. 2A and A’), consistent with a loss in apical membrane support. The thinned areas lack canalicular vesicles relative to other areas where the terminal web is still intact. These vesicles connect transiently to the canals and are believed to regulate acidification and osmotic regulation of the animal (KOLOTUEV *et al.* 2013). The lumen of the cystic canals contains visible long filaments (Fig 2B’) that have not been detected in wild-type canals or in most other *exc* mutant canals, save for mutants in the *sma-1* β_H_spectrin gene. These defects are specific to the excretory canals; electron microscopy of the intestine shows an apparently normal terminal web surrounding well-formed and normally arranged microvilli, with no cystic effects (Fig. 2C and C’).

**Figure 2.**
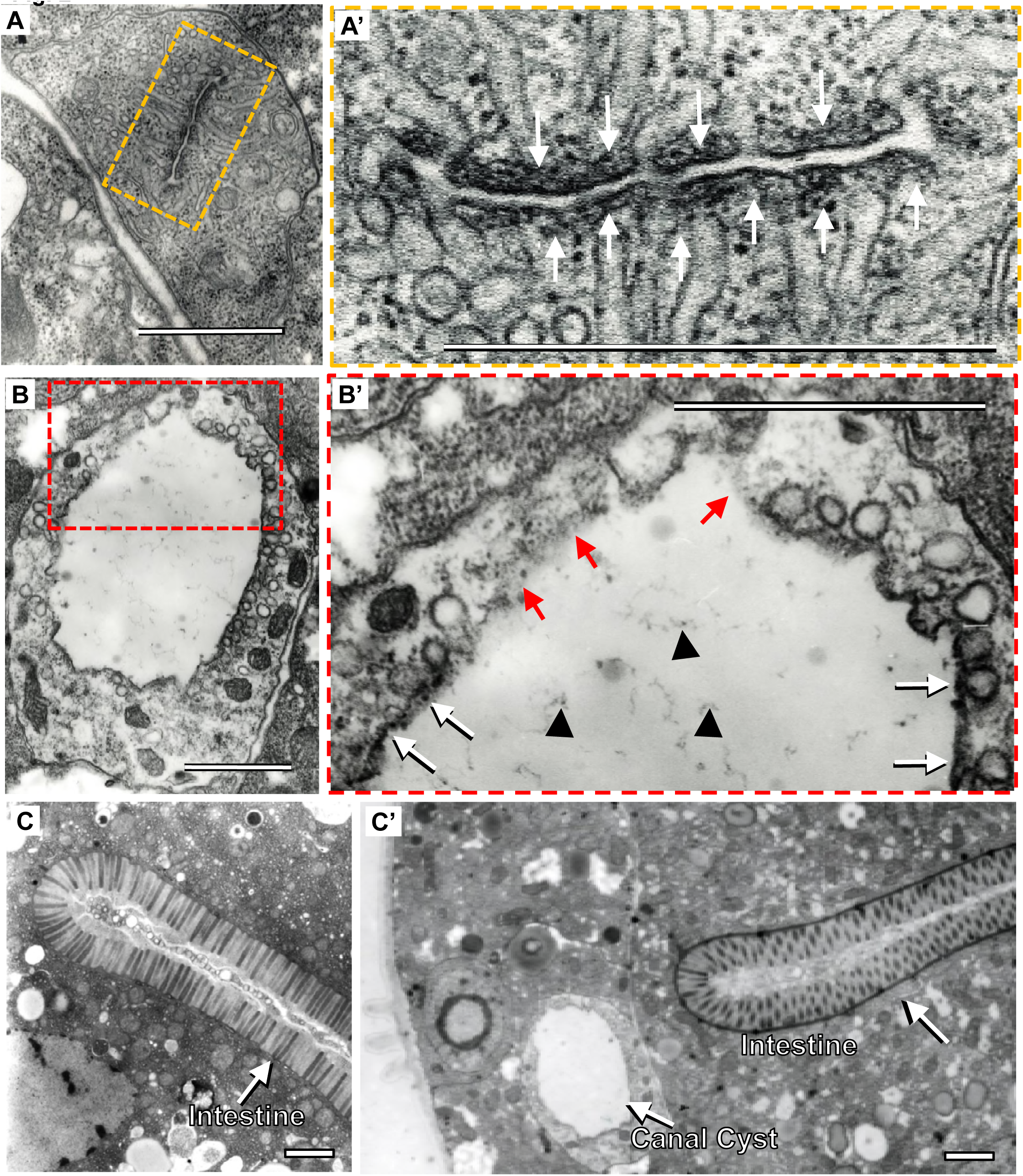
Electron microscopy of *exc-2*(*rh209*) excretory canal and intestine. Cross-sectional electron-microscopic images of wild-type (N2) and mutant (*exc-2*(*rh209*)) tissues. (A) Wild-type canal; area outlined in yellow contains lumen (connected to myriad small canaliculi) and is magnified in A’. White arrows point to the dark actin-rich terminal web surrounding the lumen on all sides. (B) *exc-2* (*rh209*) canal; area outlined in red is magnified in B’. White arrows point to thick terminal web where present; red arrows point to regions of apical membrane lacking visible terminal web. Black arrowheads indicate presumed luminal scaffold material accumulating abnormally in the lumen of multiple *exc-2* mutant alleles, but not visible in lumen of wild-type canals or other *exc* mutants. (C) Lower-magnification image of intestine in (C) wild-type and (C’) *exc-2* (*rh209*) shows intact intestine with normal arrangement of microvilli and normal basal membrane surrounded by intact terminal web surrounding entire apical surface. Mutant also shows cystic canal. Scale bars, 1µm.

Whole-genome sequencing of four *exc-2* alleles revealed that the intermediate filament gene *ifc-2* is mutated in all four strains (Fig. 3A, Supp. Fig. 2). Two of these alleles, *rh209* and *rh247*, encode nonsense mutations in exon ten and twelve, respectively. Alleles *rh90* and *rh105* include deletions of multiple coding regions that cause frameshift mutations that lead to early stop codons (Fig. 3A). In order to confirm the identity of the *exc-2* gene, a null-allele (*qp110*) in *ifc-2* was generated via CRISPR/Cas9-induced deletion; this allele harbors a deletion in the 5’ region of the gene, including the third exon of the 5’UTR along with part of first exon of the coding sequence of the gene (Fig. 3A). The *qp110* strain showed similar canal length and cyst number and size as for *exc-2*(*rh247*) (Fig. 3B and B’), and these two alleles failed to complement each other. To further confirm the identity of the *exc-2* gene, dsRNA targeting the twelfth exon of the *ifc-2* coding region was injected into wild-type worms. These worms exhibited canal defects equivalent to those in *exc-2* mutant animals (Fig. 4B and B’). As a control, injection of this dsRNA into *exc-*2 animals did not noticeably exacerbate canal defects. Finally, a rescue assay was conducted via injection, either of the ∼51kb fosmid WRM0630A_E08 containing the *ifc-2* gene (as well as *lpr-7, m6.11, pha-2, m6.4*, and a part of *rund-1*), or of the *ifc-2* gene and regulatory element (2kb upstream and 500bp downstream, total length 15.3kb) PCR-amplified from the WRM0630A_E08 fosmid, together with a GFP marker construct, into the gonad of *exc-2*(*rh247*) worms. Multiple concentrations were tested for injection (Table 2). Of the surviving progeny labeled in the canals, up to 16% showed complete rescue via fosmid injection at 12.5 ng/μl, with a full-length canal and complete absence of cysts (Fig. 3C). An additional 9% were partially rescued by fosmid, exhibiting canals shorter than wild-type, but much longer than those of *exc-2*(*rh247*) animals, with no cysts visible along the canal length. For the amplified gene at a concentration of 15 ng/μl, 19% were fully rescued plus 15% partially rescued. The relatively low rate of rescue was significantly higher than occurred through control injection of a different intermediate filament gene, *ifa-4*, 0% (Table 2). The *exc-2*(*qp110*) strain showed a similar low but significant rate of recovery (9%) when injected with the rescuing amplified *ifc-2* gene product (Table 2). The low but overall significant rescue rate for these injections is consistent with a dosage-dependent level of *exc-2* expression being necessary for wild-type canal formation. We conclude that the intermediate filament protein IFC-2 is encoded by the *exc-2* gene, and will refer here to the gene and protein by the prior name, EXC-2.

**Table 2.**
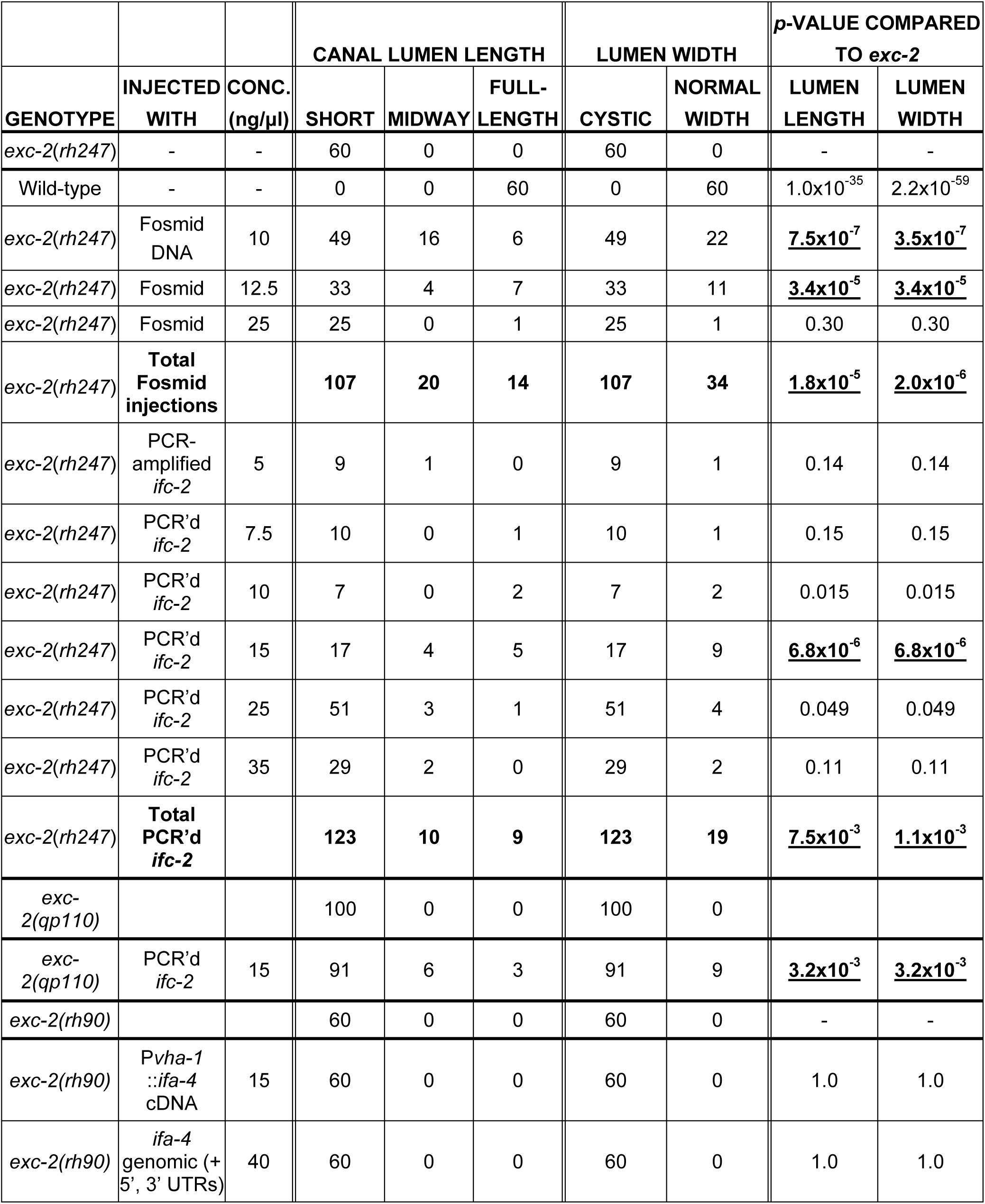
Rescue of Exc-2 phenotype by injection of *ifc-2* DNA. Phenotypes of progeny animals expressing GFP after injection into *exc-2*(*rh247*) or *exc-2*(*rh90*) hermaphrodites of the indicated constructs, as shown in Fig. 3A. Combined results for injection of Fosmid DNA at all concentrations attempted, and of PCR-amplified ifc-2 DNA at all concentrations attempted, are also included. A Fisher’s 3x2 test compared both lumen length and cyst size to canals of mutants of the appropriate genotype. Significant differences are underlined.

**Figure 3.**
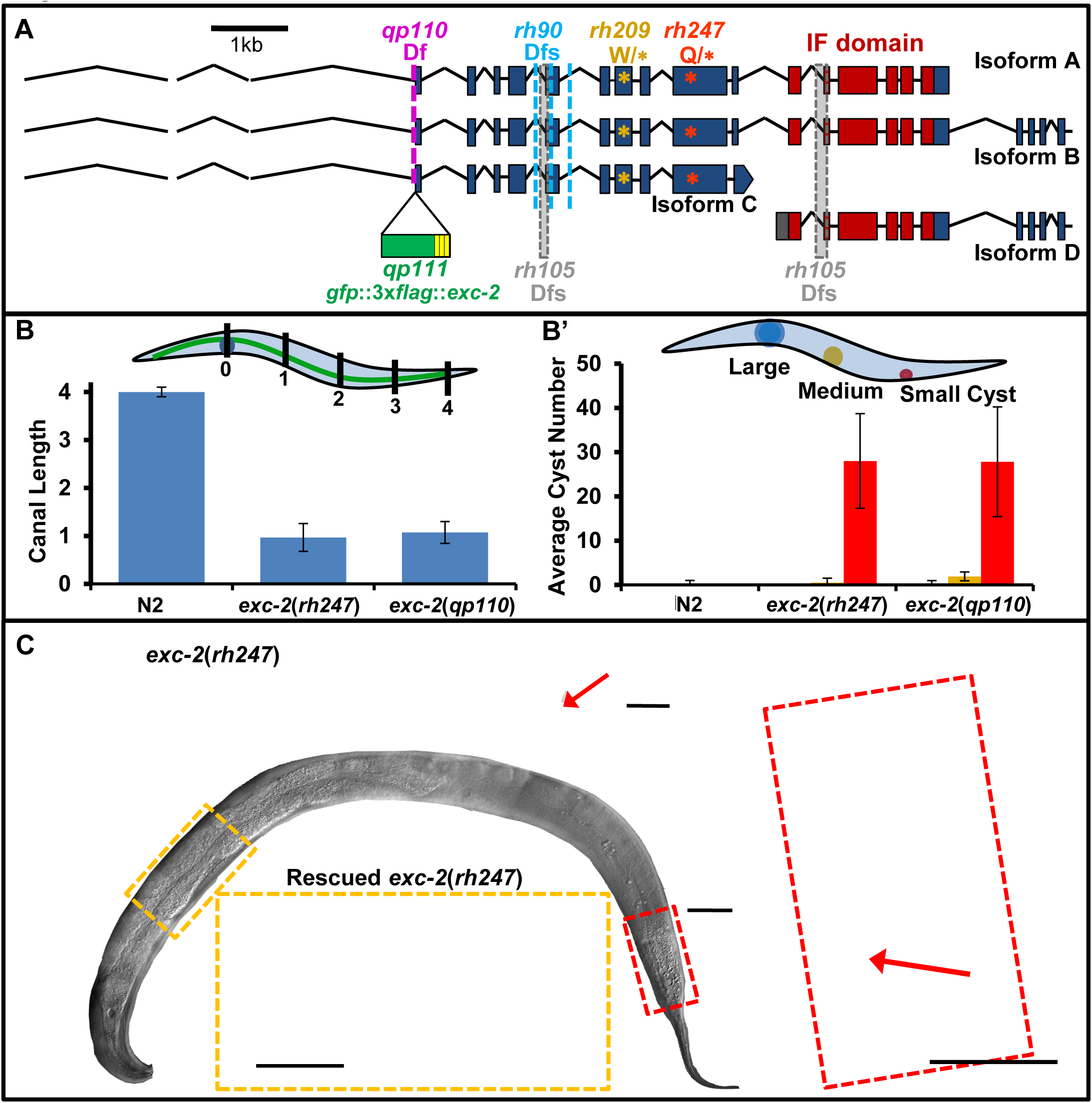
*exc-2* encodes the intermediate filament protein IFC-2. (A) Structure of *exc-2* gene (from Wormbase, release WS262). Conserved region homologous to Intermediate Filament Domain is shown in dark red. Alleles *rh209* and *rh247* contain nonsense mutations in exon ten and twelve, respectively. Alleles *rh90* and *rh105* include deletions in multiple coding regions that cause frameshift mutations, while CRISPR/Cas9-generated allele *qp110* deletes part of the promoter and 2 bases of the start codon of Isoforms A, B, and C. CRISPR/Cas9-generated allele *qp111* inserts *gfp* linked to 3 copies of FLAG-tag sequence at the start codon of isoforms A, B, and C. Bar, 1kb. (B) *exc-2*(*qp110*) mutants (CRISPR/Cas9-generated deletion at the start codon) exhibit similar canal defects to those of other *exc-2* mutants, as measured by canal length (B) and cyst size (B’); n=50. (C) Phenotypic rescue of progeny animal of *exc-2*(*rh247*) animal injected with PCR-amplified *ifc-2* gene, including 2kb upstream and 500bp downstream at 15 ng/ml. 19% of animals were completely rescued, 15% were partially rescued by this concentration of injected gene (Table 2). C’ and C’’ show magnification of boxed areas. Red arrows indicate terminus of canal. Bars, 50 µm.

**Figure 4.**
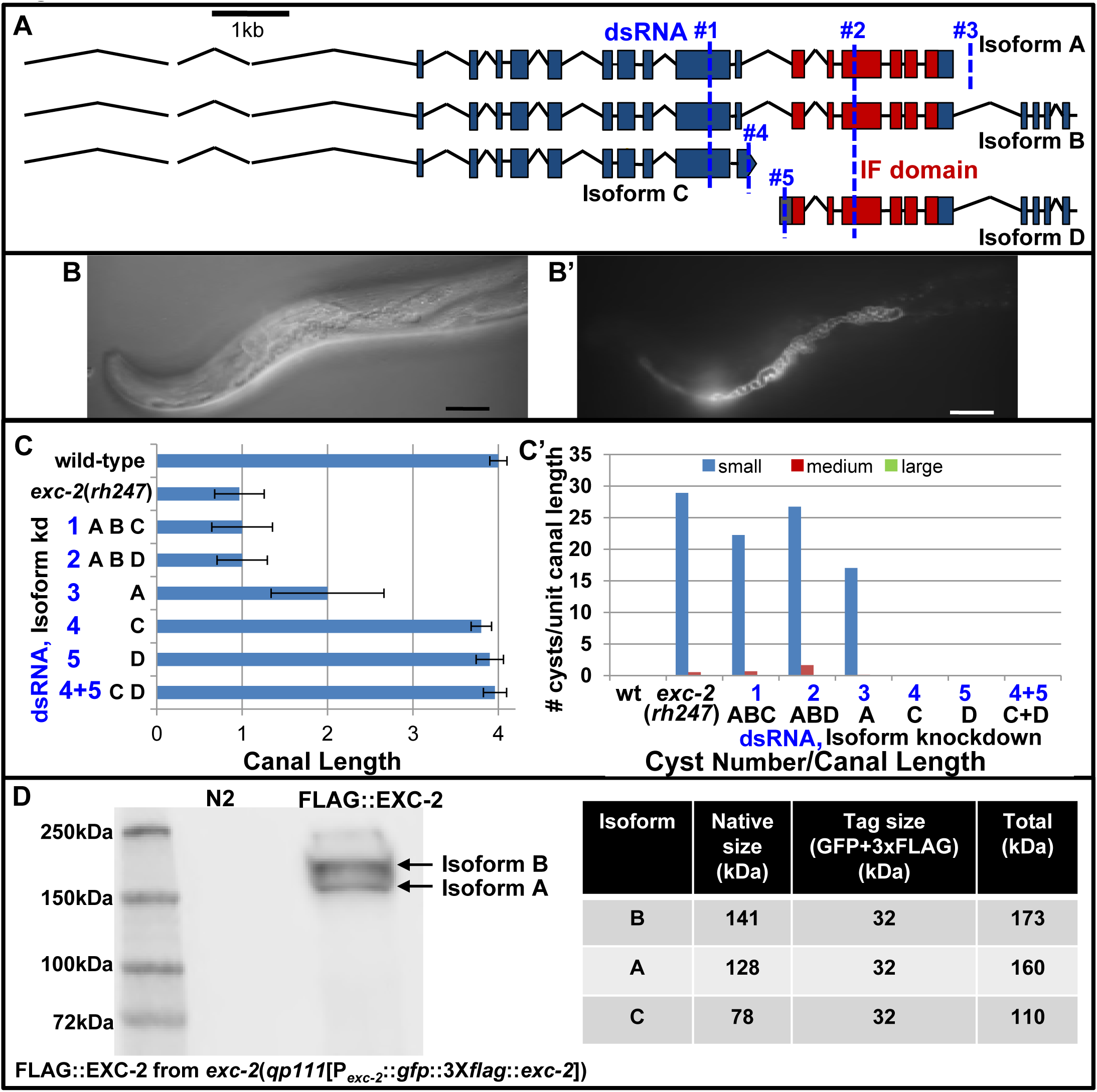
EXC-2 Isoforms A and B maintain canal structure. (A) Schematic diagram of isoforms of *exc-2* (from Wormbase release 262) showing positions of RNAi-targeted regions (blue dashed lines). Bar, 1 kb. (B) DIC and (B’) fluorescence images for progeny of injected worms with dsRNA #1 targeting the twelfth exon of *ifc-2*. Animals presented canals as short and cystic as those of *exc-2*(*rh247*) animals. (C) Measurements of canal length (C) and cyst number (C’) in different isoform-specific RNAi knockdown animals. (n>50 each). (D) Protein expression of N-terminal FLAG-tagged EXC-2. Western blot shows two predominant bands corresponding to predicted sizes of isoforms A and B.

In order to identify the isoforms of EXC-2 that are responsible for the excretory canal phenotype, we directly injected dsRNA into wild-type worms (synthesized and transcribed *in vitro*) targeted to specific *exc-2* isoforms (Fig. 4A). The canals in progeny animals were then evaluated with regard to canal length, and cyst number and size (Fig. 4C and C’’, Table 3). The first dsRNA (#1 in Fig. 4A) targeted the twelfth exon, which knocks down isoforms A, B, and C. These worms showed a canal phenotype similar to that of *exc-2*-null strains in both length and cyst size (Fig. 4B). dsRNA #2 targets the sixteenth exon to knock down isoforms A, B, and D, and this knockdown also resulted in a phenotype similar to that of *exc-2* mutants. As antibodies to Isoform D bind to the intestine (KARABINOS *et al.* 2001), and long-term knockdown of Isoform D has been reported to affect intestinal structure (KARABINOS *et al.* 2004), we also examined this organ in progeny animals, but saw no intestinal effects in progeny, even in animals exhibiting strongly cystic canals (Supp. Fig. 3). dsRNA #3 targeted the 3’UTR solely in isoform A, in exon nineteen. These worms showed a milder phenotype in which the canal length reached the vulva midway along the length of the animal, and displayed cysts, but smaller than those of *exc-2* knockout animals. dsRNA #4 targeted the coding sequence that is uniquely transcribed at the end of isoform C. This knockdown had no effect on canal length, and no cysts were formed. Finally, dsRNA #5 targeted the 5’UTR of isoform D. Canal length was as long as in wild-type animals, and no cysts were observed, and again, no intestinal effects were observed. To confirm the lack of effects of knockdown of isoforms C and D, we injected a mixture of dsRNA(s) targeting both isoforms. These worms showed no deleterious effects on the canals. These results indicate that both isoforms A and B are needed for EXC-2 function in canal formation.

**Table 3.** Effects of isoform-specific RNAi-knockdown of *exc-2*. Phenotypes of progeny animals expressing GFP after injection into hermaphrodites of the indicated dsRNAs, as shown in Fig. 4A. A Fisher’s 3x2 test compared both lumen length and cyst size to canals of wild-type animals. Significant differences are underlined.

|  |  | CANAL LUMEN LENGTH |  |  | LUMEN CYST SIZE |  |  | *p*-VALUE<br>COMPARED TO<br>WILD-TYPE |  |
| --- | --- | --- | --- | --- | --- | --- | --- | --- | --- |
| WILD TYPE<br>INJECTED<br>WITH RNAi# | ISOFORMS<br>TARGETED | SHORT | MIDWAY | FULL-<br>LENGTH | LARGE+<br>MEDIUM<br>CYSTS | SMALL<br>CYSTS | NO<br>CYSTS | LUMEN<br>LENGTH | CYST<br>SIZE |
| None<br>(wild type) |  | 0 | 0 | 60 | 0 | 0 | 60 | - | - |
| 1 | A, B, D | 42 | 16 | 0 | 9 | 48 | 0 | <u>$4.2 \times 10^{-35}$</u> | <u>$8.5 \times 10^{-35}$</u> |
| 2 | A, B, C | 49 | 2 | 0 | 7 | 44 | 0 | <u>$7.3 \times 10^{-33}$</u> | <u>$7.3 \times 10^{-33}$</u> |
| 3 | A | 9 | 42 | 1 | 0 | 49 | 0 | <u>$2.07 \times 10^{-31}$</u> | <u>$3.5 \times 10^{-32}$</u> |
| 4 | C | 0 | 1 | 28 | 0 | 0 | 25 | 0.326 | 1.0 |
| 5 | D | 0 | 1 | 61 | 0 | 0 | 62 | 0.999 | 1.0 |
| 4 + 5 | C, D | 0 | 0 | 50 | 0 | 0 | 50 | 1.0 | 1.0 |
| 3 + 4 + 5 | A, C, D | 0 | 26 | 7 | 0 | 33 | 0 | <u>$5.4 \times 10^{-17}$</u> | <u>$6.2 \times 10^{-26}$</u> |

In order to validate the dsRNA data, we looked at total protein levels of isoforms A, B, and C in strain BK531 (“tagged *exc-2* strain”) in which *exc-2* was modified via CRISPR/Cas9 to place *gfp* and three copies of a *flag* tag just upstream of the starting AUG codon of these three isoforms. The western blot (Fig. 4D) showed only two large isoforms in the worm lysate sample, corresponding to the size of isoforms A and B. It should be noted that the anti-FLAG antibody cannot detect isoform D; the result is still in agreement with the knockdown results, however. We conclude that both isoforms A and B are needed for EXC-2 function in the excretory canals.

Examination of strain BK531 (tagged *exc-2*) showed the expression pattern of the *exc-2* gene (Fig. 5). Labeled EXC-2 is located at the lumen of the excretory canal, in the corpus, posterior bulb, and pharyngeal-intestinal valve (PIV) of the pharynx, as well as in the uterine seam (UTSE) and intestinal-rectal valve (VIR). The subcellular location of EXC-2 in these tubes was compared to that of exogenously expressed cytosolic mCherry (Fig. 6). Labeled EXC-2 is located apical to canal cytoplasm (Fig. 6A-A’’), as determined via cross-sectional fluorescence intensity measurements (Fig. 6A’’’). This result was confirmed by evaluating the subcellular location of EXC-2 relative to a known apical membrane protein, ERM-1 (GÖBEL *et al.* 2004). The results demonstrate that EXC-2 and ERM-1 show overlapping expression at the canal apical (luminal) membrane (Fig. 6B-B’’’).

**Figure 5.**
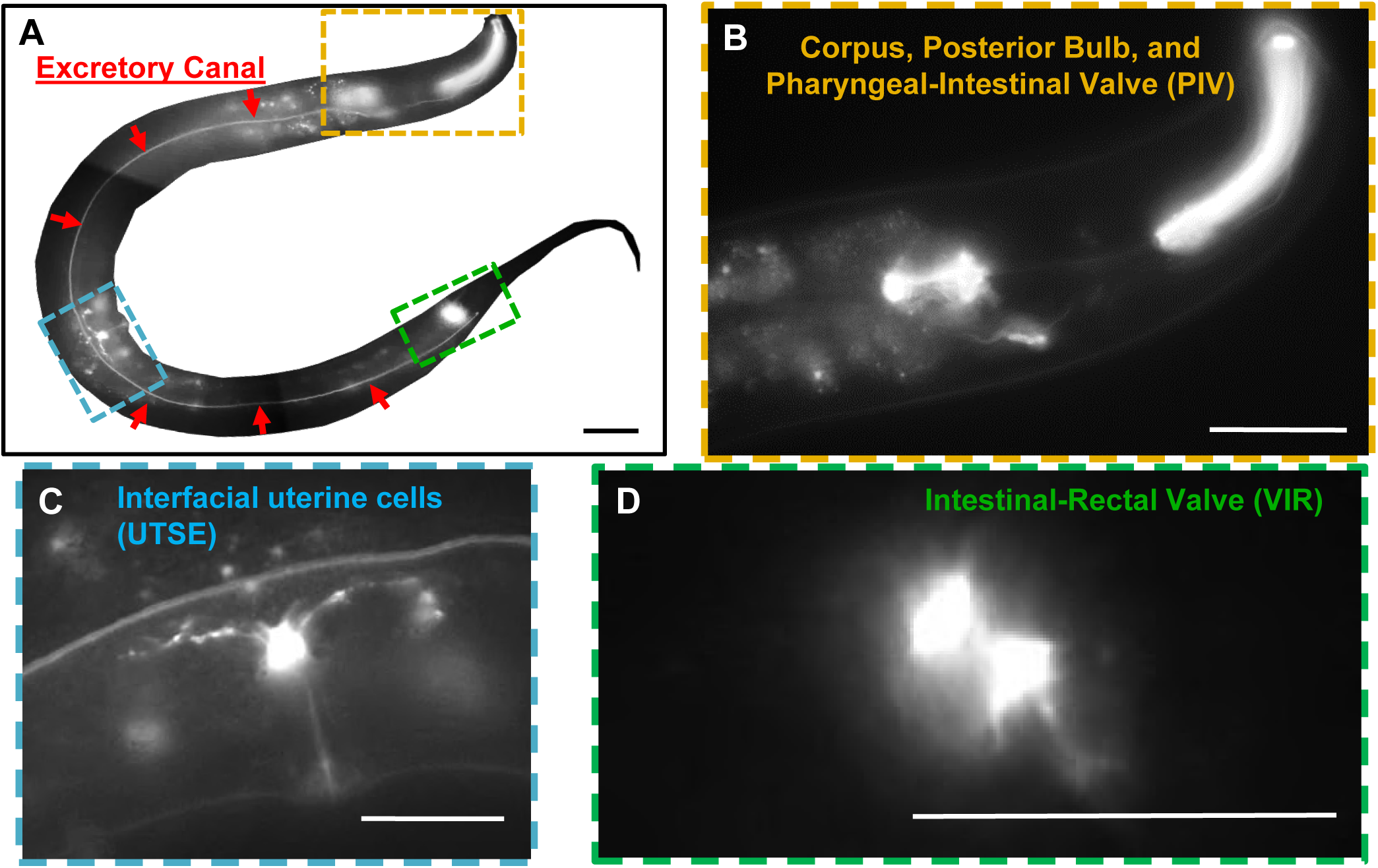
EXC-2 is expressed in multiple epithelial cells. A) A CRISPR/Cas9 knock-in of *gfp* at the N-terminus of *exc-2* is expressed in four tissues: 1) The excretory canals (A,C); 2) The pharyngeal corpus, posterior bulb, and pharyngeal-intestinal valve (PIV) (B); 3) The interfacial uterine cell (UTSE) (C), and; 4) The intestinal-rectal valve (VIR) (D) Bar, 50µm.

**Figure 6.**
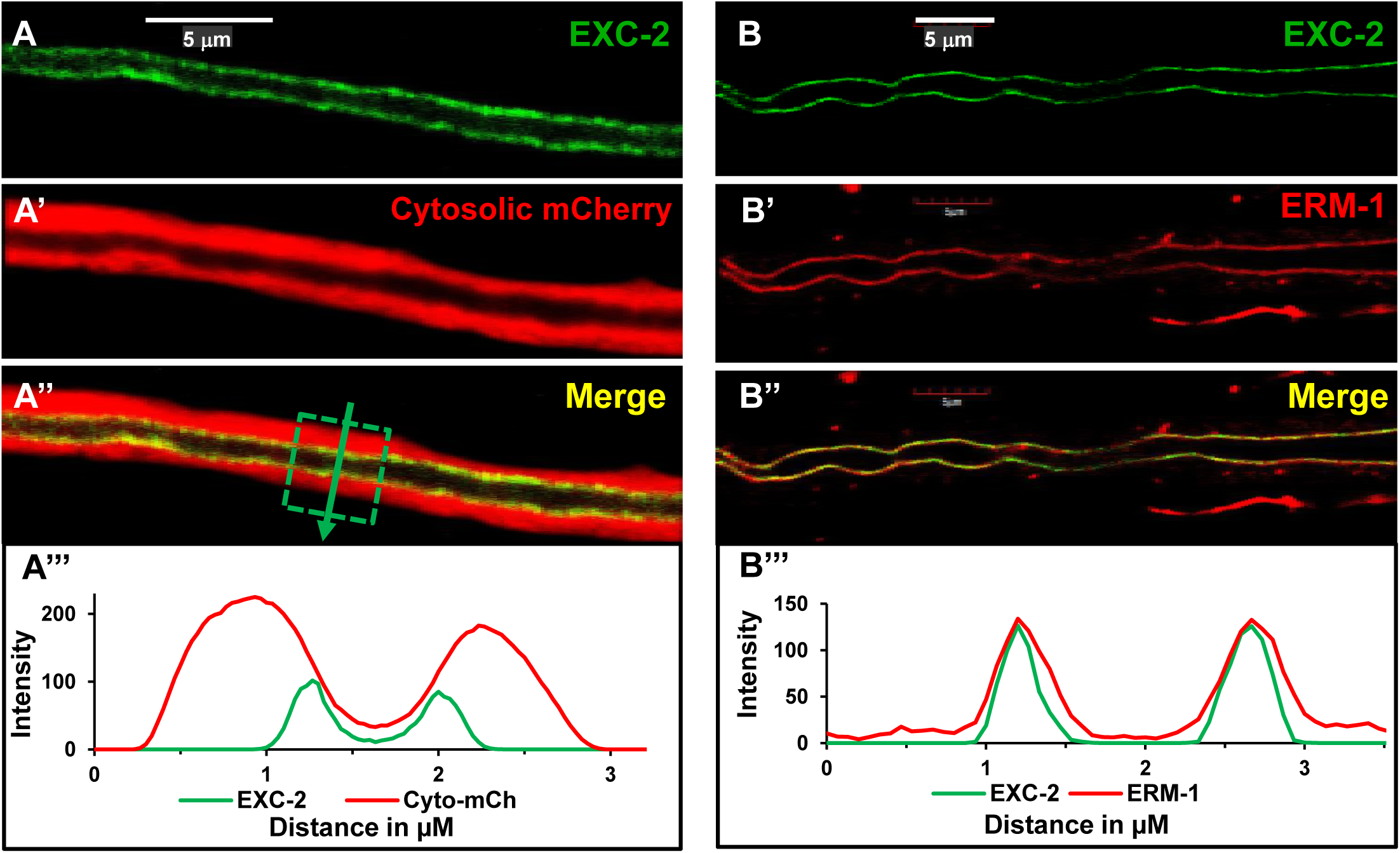
EXC-2 is expressed at the apical membrane of the excretory canals. (A) Apical position of EXC-2 (GFP) relative to (A’) canal cytosol labelled with mCherry; (A’’) merged panel. (A’’’) Graph of fluorophore intensity for each color within canal section (box in A’’) along length of arrow. (B) Coincident position of EXC-2 (GFP); with (B’) apical epithelial marker ERM-1 labeled with mCherry; (B’’) merged panel. (B””) Graph of fluorophore intensity measurement within canal section boxed in B’’. Bars, 5µm.

IFA-4 is the third intermediate filament gene that is highly enriched in the canal (SPENCER *et al.* 2011). We investigated whether IFA-4 plays a role in canal maintenance similar to that of EXC-2 and IFB-1 (WOO *et al.* 2004). The *ifa-4*(*ok1734*) strain from the CGC carries a deletion of the *ifa-4* coding sequence, and exhibits a canal phenotype similar to that of *exc-2* mutants (Fig. 7A-A’’). Injection of dsRNA specific to *ifa-4* phenocopies the short cystic canals of the *ifa-4* deletion strain (Fig. 7B-B’’). Strain BK532 (“tagged *ifa-4* strain”) was created via CRISPR/Cas9 to tag *ifa-4* with *mKate2* and three *myc*-tags at the 5’ end of the coding region (Fig. 7C). 40% of these animals exhibited wild-type morphology, while in 60% of these animals (which tended to have brighter expression) the canals were slightly shortened (to length ∼3.5), but did not exhibit any cysts. IFA-4::mKate2 showed expression in several of the same locations as *exc-2*: the excretory canals, the pharyngeal-intestinal valve, and the intestinal-rectal valve. IFA-4 is additionally located in the spermathecal-uterine valve and cells in the vulva, including the uterine muscles and two neurons in the tail (Fig. 7C). The IFA-4 protein was also found in the gut of dauer-stage but not well-fed worms, and appeared as well in the tips of the rays and neurons of the male tail (Supp. Fig. 4). An overexpressing translational construct (*P*_*vha-1*_*::ifa-4::gfp*) rescued the mutant canal phenotype of the *ifa-4*(*ok1734*) deletion mutant strain, and showed subcellular expression of this protein at the apical membrane of the canal, though with large inclusions protruding deeper into the cytosol (Fig. 8A-A’’’). The 60% of BK532 (tagged *ifa-4*) animals that exhibited shortened canals also contained these subcellular inclusions. Finally, subcellular collocation of IFA-4 and EXC-2 at the canal apical membrane was confirmed in strain BK533 carrying CRISPR/Cas9-tagged insertions both of *gfp* into the 5’ end of *exc-2* and of *mKate2* into the 5’ end of *ifa-4* (Fig. 8B-B’’’).

**Figure 7.**
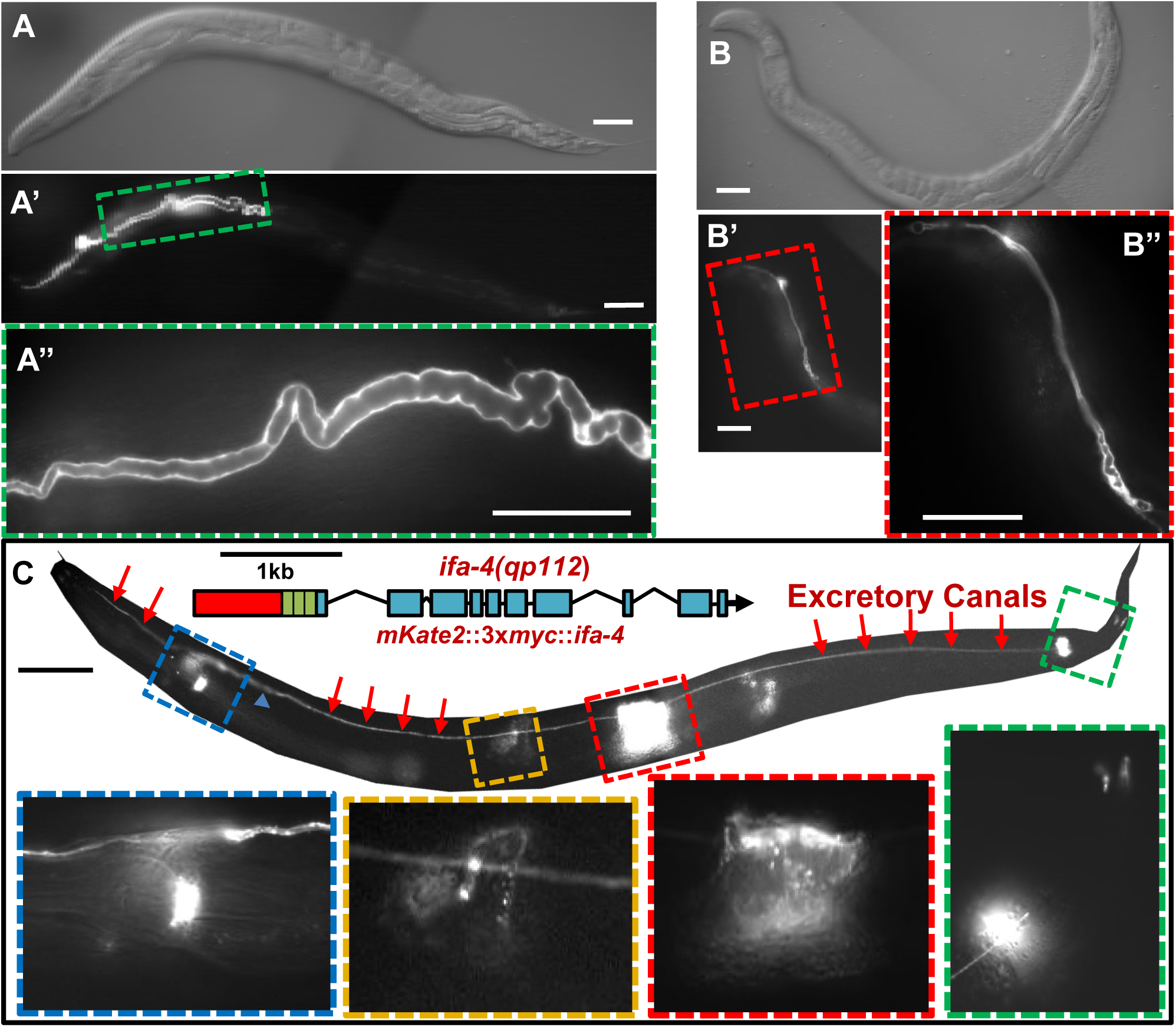
IFA-4 effects canal maintenance. (A) DIC and (A’, A’’) fluorescence micrographs of excretory canal of strain BK535 (*ifa-4* ko; canal marker). Boxed area of (A’) magnified in (A’’). (B) DIC and (B’) fluorescence micrographs for progeny of animal injected with dsRNA targeting the seventh exon of *ifa-4*. Animals exhibited short, cystic canals. Boxed area of (B’) magnified in (B’’). (C) Diagram of CRISPR/Cas9 knock-in of *mKate2* at the N-terminus of *ifa-4*. *ifa-4* is expressed in five tissues: 1) The excretory canals (red arrows); 2) Pharyngeal-intestinal valve (PIV) (blue box); 3) The spermathecal-uterine valve (yellow box); 4) Uterine muscles (red box), and; 4) The intestinal-rectal valve and neurons at the tip of the tail (green box). Boxed areas are below main image, outlined in boxes of the corresponding color. All bars, 50µm.

**Figure 8.**
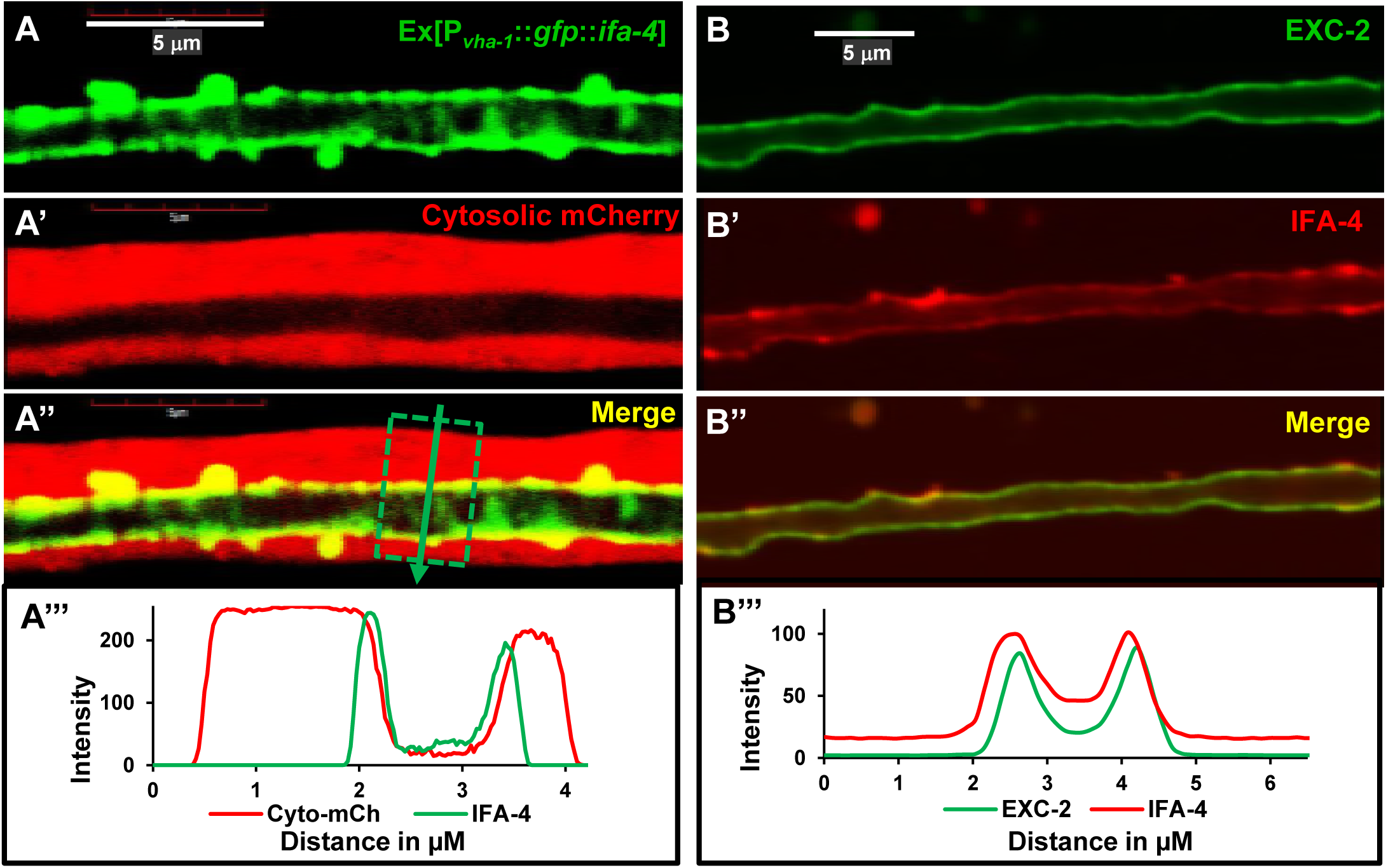
IFA-4 is co-localized with EXC-2 to the apical membrane. (A). Fluorescence micrographs of injected (40ng/µl) translational construct (P_*vha-1*_::*ifa-4*::*gfp*) ectopically expressed in *ifa-4*(*ok1734*) deletion mutant, which fully rescued canal morphology. (A’) Cytosolic mCherry marker co-expressed in the canal; (A’’) Merged image. (A’’’) Graph of fluorophore intensity for each color within canal section (box in A’’) along length of arrow. (B) Fluorescence micrographs of CRISPR/Cas9-modified strains labelling *exc-2* and *ifa-4*. (B) shows GFP::EXC-2 expression, (B’) mKate2::IFA-4 expression, (B’’) merged image. (B’’’) Graph of fluorophore intensity for each color within canal section. Bars, 5µm.

Several *exc* genes (*exc-1, exc-5, exc-9*) with knockout canal phenotypes of large cysts also exhibit characteristic overexpression phenotypes that rescue the canal lumen diameter, while shortening canal length, and show epistatic genetic interactions (TONG and BUECHNER 2008; MATTINGLY and BUECHNER 2011; GRUSSENDORF *et al.* 2016). We therefore looked at overexpression phenotypes of *exc-2* and *ifa-4*. PCR-amplified *exc-2* that rescued *exc-2* mutants (Fig. 3C) was injected (together with a fluorescent canal marker) into N2 wild-type worms (Fig. 9A). All progeny showing the injection marker also exhibited substantially shortened canals extending only to the vulva (average canal length of 2.1 (n=14)), but with small or no cysts. We created a similar construct of PCR-amplified *ifa-4* cDNA, linked to gfp, under the control of a strong canal-specific promoter. After injection into wild-type worms, progeny expressing GFP (Fig. 9A’). showed shortened canals, extending just beyond the vulva (length 2.5, n=30). The similarity of overexpression phenotype between the two intermediate filament genes is consistent with these proteins performing similar functions.

**Figure 9.**
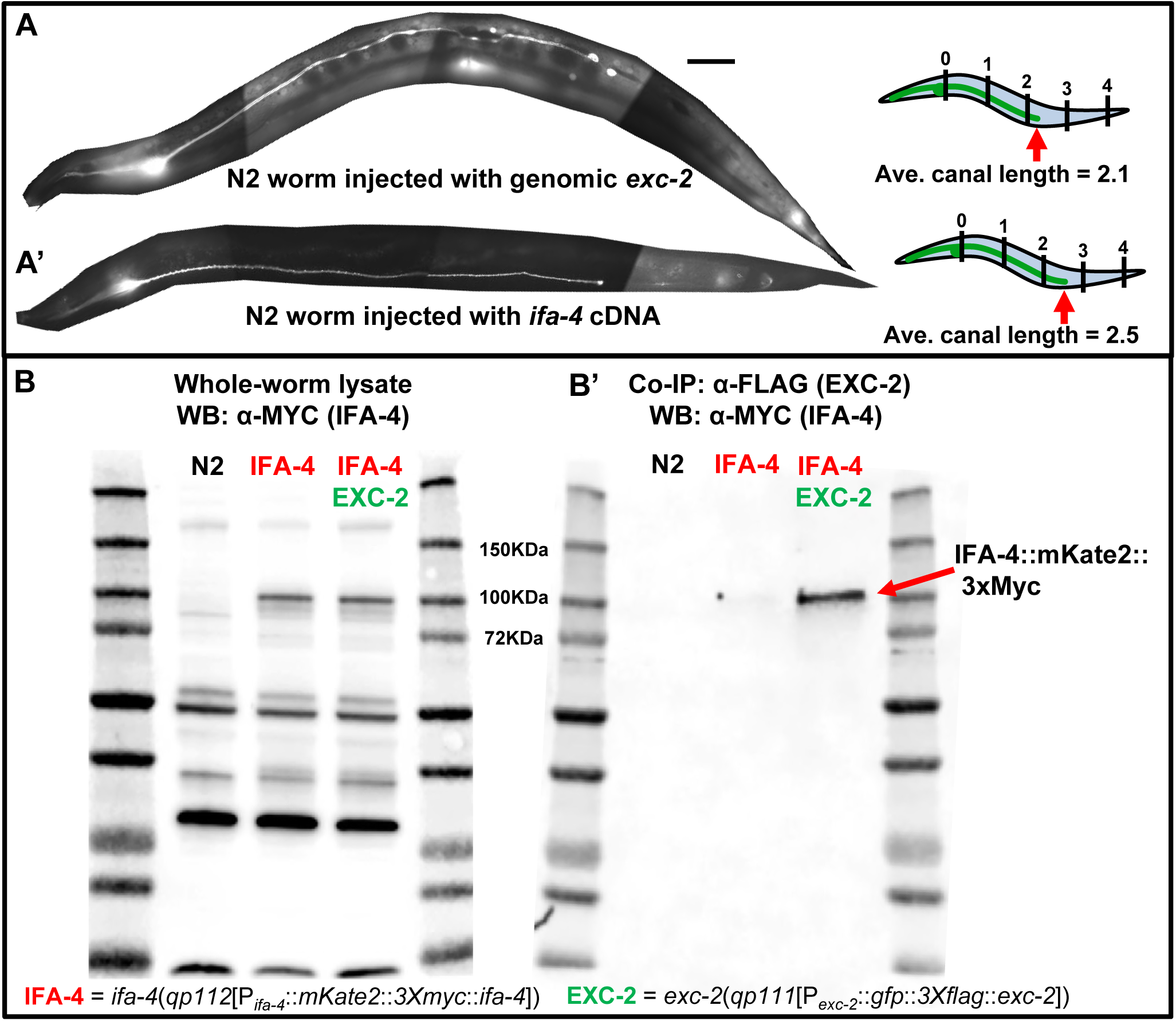
EXC-2 and IFA-4 interact directly. A) Representative N2 worms injected with (A) *exc-2* genomic construct (including 2kb upstream and 500bp downstream of coding region) or (B) *ifa-4* cDNA construct under control of strong canal promoter *vha-1* to overexpress these proteins. Both lines exhibited shortened canal lumen of wild-type diameter with no cysts. For *exc-2* overexpression, length=2.1 (n=14); for *ifa-4* overexpression, length=2.5 (n=30). Bars, 50µm. B) Western blot of whole-worm lysates probed with antibody to MYC. (B’) Western blot of lysates purified via anti-FLAG magnetic beads, which bind to FLAG-tagged EXC-2. Lysates are from wild-type animals (N2, left), animals expressing Myc-tagged IFA-4 (middle), and animals expressing both Myc-tagged IFA-4 and FLAG-tagged EXC-2 (right); lanes on blots are flanked by size marker lanes. Red arrow in blot on right indicates size of IFA-4 band at ∼100kDa.

A co-immunoprecipitation assay was conducted to test whether EXC-2 and IFA-4 bind each other directly (Fig. 9B and B’). Protein lysates were prepared from three worm strains: Wild-type (N2); BK532 (tagged *ifa-4*), and BK533 (tagged *exc-2* & *ifa-4*). Tagged IFA-4 could be detected in blots of whole-worm lysates (Fig. 9B), but was only detectable in αFLAG pulldowns when tagged EXC-2 was present (Fig. 9B’), which indicates that the two proteins bind to each other.

The interaction of EXC-2 with IFA-4 led us to investigate whether the apical localization of EXC-2 in the canals depends on the function of the other two intermediate filaments. Strain RB1483, carrying the *ifa-4*(*ok1734*) deletion, was crossed to strain BK531 (tagged *exc-2*) (Fig. 10A-A’’’’), then injected with a cytosolic mCherry marker construct. The subcellular location of EXC-2 at the apical membrane of the canal was unchanged in these *ifa-4* knockout mutants.

**Figure 10.**
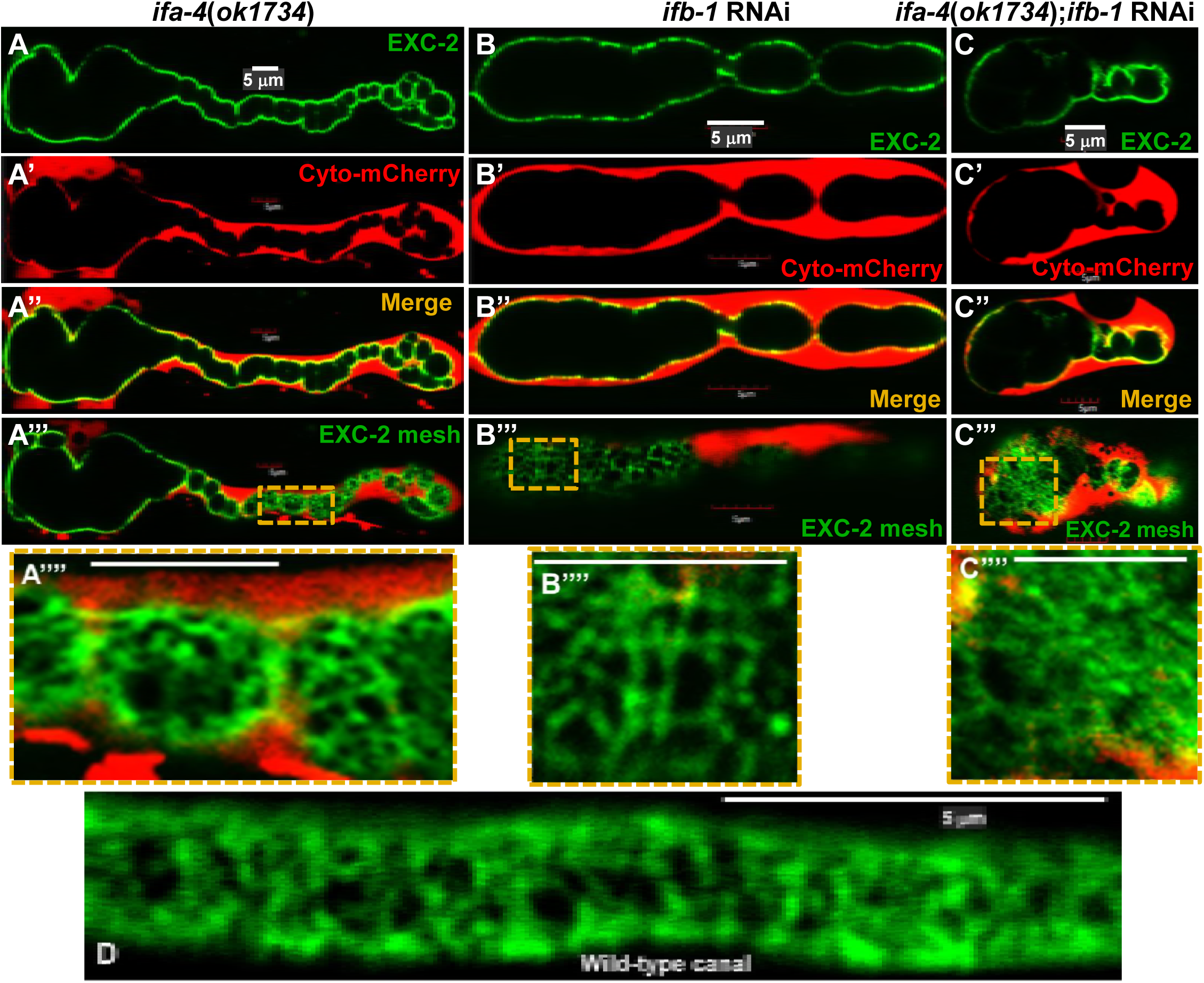
EXC-2 apical location is independent of IFB-1 and IFA-4 function. Fluorescence micrographs of GFP-tagged EXC-2 in: (A) *ifa-4* knockout strain (*ok1734*); (B) animal exhibiting canal-specific RNAi-knockdown of *ifb-1*; (C) RNAi-knockdown of *ifb-1* in *ifa-4* knockdown strain. All animals express cytosolic mCherry marker. (A,B,C) GFP::EXC-2 fluorescence; (A’,B’,C’) Cytosolic mCherry fluorescence; (A’’,B’’,C’’) Merged images. (A’’’, B’’’, C’’’) Plane of focus higher in Z-plane, to observe merged fluorescence of GFP::EXC-2at apical surface of swollen cystic areas of canals flattened against hypoderm.. Boxed areas are magnified in panels A’’’’, B’’’’, and C’’’’. (D) Fluorescence of GFP::EXC-2 at the narrower surface next to hypoderm in wild-type canals. All bars, 5µm.

The apical location of EXC-2 was similarly tested in the absence of IFB-1. Due to embryonic lethality of *ifb-1* null mutants (due to hypodermal defects (WOO *et al.* 2004)), a construct expressing dsRNA to *ifb-1* under a canal-specific promoter was injected together with cytosolic mCherry marker construct into strain BK531 (tagged *exc-2*) (Fig. 10B-B’’’’). These animals exhibited cystic canals consistent with knockout of IFB-1, and EXC-2 retained its apical location in the presence of this *ifb-1* knockdown. Finally, we investigated whether the apical location of EXC-2 depends on the presence of either one of the other intermediate filaments, IFA-4 or IFB-1 (Fig. 10C-C’’’’). Injection of the same canal-specific dsRNA construct against *ifb-1* along with cytosolic mCherry marker construct into the *ifa-4*(*ok1734*) mutant strain and crossed to BK531 (tagged *exc-2*) resulted in the same location for EXC-2. The double knockout was sickly, with lethality higher than for either *ifb-1* knockdown or *ifa-4* mutation alone (n=97, Supp. Fig. 5), consistent with an additive function of IFA-4 and IFB-1. Death in these animals occurred throughout mid-larval stages, much later than the two-fold embryonic muscle failure and death of *ifb-1* null mutants (WOO *et al.* 2004), which suggests a general inability of canals to function effectively in the combined abseence of these two intermediate filament proteins. (BUECHNER *et al.* 1999; HAHN-WINDGASSEN and VAN GILST 2009).

Finally, the subcellular expression pattern of EXC-2 formed a mesh-like network on the apical surface of the canals, both in wild-type canals (Fig. 10D) and in animals lacking IFA-4, IFB-1, or both (Figs. 10A’’’’, B’’’’, and C’’’’). This meshwork is more easily observed on cysts, where a larger surface area is in the focal plane, than in the narrow canals of wild-type animals, but the size and general arrangement of filaments appears similar in all cases, which indicates that EXC-2 filaments can be localized to the apical membrane even without the function of the other two canal intermediate filament proteins.

## DISCUSSION

### ***exc-2* encodes for the intermediate filament IFC-2**

This study has used multiple methods to confirm the identity of the *exc-2* gene: 1) Whole-genome sequencing of four alleles; 2) Injection of dsRNA to knock down multiple isoforms of the gene; 3) Rescue of a null allele by both fosmid and PCR-amplified DNA; 4) CRISPR/Cas9-mediated tagging of the *exc-2* gene showing the tissue and subcellular location of the encoded protein at the apical surface of the tissue affected by mutation; 5) Generating a new allele of the gene, via CRISPR/Cas9-induced deletion, which caused the same phenotype; 6) Noncomplementation between the CRISPR-induced deletion allele and a previous allele, and; 7) Pulldown of the expressed tagged EXC-2 protein to another intermediate filament that is expressed at the same location in the excretory canals, and has the same genetic effects on the excretory canals.

This cloning result is surprising, since it contradicts multiple previous studies that found IFC-2 not in the excretory canals, but primarily in desmosomes of the *C. elegans* intestine (KARABINOS *et al.* 2002; KARABINOS *et al.* 2004; HÜSKEN *et al.* 2008; CARBERRY *et al.* 2012; COCH and LEUBE 2016; GEISLER *et al.* 2016), though expression of one fluorescent construct showed expression at the intestinal apical surface (COCH and LEUBE 2016). These previous studies assumed Isoform D to be the full-length IFC-2, since this isoform is about the same size as other *C. elegans* intermediate filament proteins, and some confusion of RNA-sequencing results presented on earlier versions of Wormbase (e.g. Wormbase.org, release WS180) listed up to eight splice forms and suggested the possibility that the first 13 exons (i.e. Isoform C) comprised a separate gene. More extensive RNA sequencing since that time (summarized on current Wormbase release WS262) shows that although Isoform C and Isoform D do not overlap, both are transcripts from different parts of the same gene, and that the large Isoform A in fact comprises the entire 24 exons of the gene. Western blot of FLAG-labelled EXC-2 protein in the present study (Fig. 4D) matches the current predicted protein sizes on Wormbase. We also note that the pioneering comprehensive study of *C. elegans* intermediate filament genes (DODEMONT *et al.* 1994) reported eight intermediate filament genes, including *ifc-2*, and presents a Northern blot (Dodemont *et al.* ’94, Fig. 4) showing two much larger transcripts that correspond in size to the Isoform A and B mRNAs. Subsequent work on IFC-2 from the Weber, Leube, and Karabinos labs (KARABINOS *et al.* 2001; KARABINOS *et al.* 2002; KARABINOS *et al.* 2003; KARABINOS *et al.* 2004) created and used a polyclonal antibody to a fragment of the conserved intermediate filament domain of IFC-2; this antibody bound a single ∼55kDa protein on blots (KARABINOS *et al.* 2003), and further studies showed this antibody binding specifically to proteins in the intestine (HÜSKEN *et al.* 2008), as long-term treatment with RNAi to *ifc-2* removed intestinal staining but not staining of other tissues. Our CRISPR-tagged GFP was inserted at the first codon of Isoforms A, B, and C, and so cannot show expression of isoform D. It is therefore possible that Isoform D is expressed in the intestine, while the other isoforms are expressed in the excretory canals and other tissues shown here. Since Isoforms A, B, and D all include the conserved intermediate filament domain, however, we cannot explain why previous studies did not find expression within the excretory canals.

Previous studies (KARABINOS *et al.* 2003; KARABINOS *et al.* 2004) found that in worms fed RNAi specific to the intermediate filament domain over the course of three generations, the animal intestines slowly acquired gross morphological effects, although they saw that at the ultrastructural level, the microvilli and terminal web appeared intact. We directly microinjected dsRNAs specific to multiple isoforms into the gonad, and found that in all cases, progeny intestines were unaffected, while the excretory canals uniformly were strongly cystic, matching the phenotype of all five of our *exc-2* mutant alleles. In particular, this result obtained for dsRNA #2 (Fig. 4A), a knockdown of the conserved intermediate filament domain of Isoforms A, B, and D. Finally, the *rh105* allele has deletions in the conserved domain of these three isoforms (Fig. 3A), and exhibits large canal cysts with no intestinal defects (Supp. Fig. 1). In summary, our results strongly suggest that, while Isoform D may be expressed in the intestine, the major locus of function of the EXC-2/IFC-2 intermediate filament isoforms is in the excretory canals.

### EXC-2, IFB-1, and IFA-4 maintain tubular morphology at the apical membrane

At 12.7kb, the *exc-2* gene is easily the largest IF gene in *C. elegans*, which helps explain the relatively high frequency of alleles of this gene identified in genetic screens for cystic canal mutants (BUECHNER *et al.* 1999). The two large Isoforms A and B of this protein had clear effects within the canals as seen by mutations at multiple sites and effects of dsRNA to all areas of the gene. No effects were noticed from knockdown solely of Isoform C or Isoform D.

A map of gene expression for specific tissues of *C. elegans* (SPENCER *et al.* 2011) examined expression in the excretory canal cell, and in addition to *exc-2*, found two other highly expressed intermediate filament genes, *ifb-1* and *ifa-4*, confirming earlier studies of intermediate filament expression (KARABINOS *et al.* 2003). Knockdown of *ifb-1* has previously been found to cause formation of cysts in the canal, as well as surprising breaks in canal cytoplasm during cell outgrowth (KARABINOS *et al.* 2003; WOO *et al.* 2004; KOLOTUEV *et al.* 2013). The present study shows that IFA-4 also is necessary to maintain canal morphology. Intermediate filaments form homo- and hetero-polymers to carry out their functions (ZUELA and GRUENBAUM 2016). While IFA-4, IFB-1, and EXC-2 are all expressed at the apical (luminal) surface of the excretory canals, they each have varied expression (and presumably function) in other tissues, both overlapping and non-overlapping. IFB-1, in particular, has an essential role in embryonic muscle attachment and hypodermal cell elongation (WOO *et al.* 2004), tissues where IFA-4 and EXC-2 are not expressed. Future studies of these tissues may find other functions for these intermediate filament proteins. For example, as stretching of dissected intestines can be measured (JAHNEL *et al.* 2016), it may be possible to determine the role of IFA-4 in intestinal integrity during dauer formation, as compared to intestines in other stages where IFA-4 is not expressed.

Ultrastructural analysis of *exc-2* mutants found visible fibrous material in the lumen (Fig. 2B’), which has also been seen only in other excretory system mutants affecting apical cytoskeleton proteins: *sma-1* (encoding β_H_spectrin, *erm-1* (ezrin-moesin-radixin homologue), and the excretory duct cell gene *let-653* (mucin) (MCKEOWN *et al.* 1998; BUECHNER *et al.* 1999; KHAN *et al.* 2013; GILL *et al.* 2016). Future studies may show whether the fibrous material represents proteins or other material normally anchored to the membrane directly or indirectly by these cytoskeletal proteins.

### Three intermediate filament proteins support the canal apical membrane

The three intermediate-filament proteins EXC-2, IFA-4, and IFB-1 are collocated at the apical membrane of the canal, have similar knockdown effects, and bind to each other in pulldown assays (Fig. 9B) and (KARABINOS *et al.* 2003)). The ratio between these proteins affects their function, as overexpression of either IFC-2 or IFA-4 allows formation of small cysts in short canals (Fig. 9A and A’).

EXC-2 forms homo- and heterodimers in *in vitro* studies (KARABINOS *et al.* 2017), which may be necessary in order to form a strong meshwork at the canal apical surface, as seen for lamins and other intermediate filament proteins at cell membranes. Lamin, for example, is bound directly to the membrane to form such structures at the inner nuclear membrane (FAWCETT 1966; DECHAT *et al.* 2008; DE LEEUW *et al.* 2017), through farnesylation of the CAAX domain at the lamin C-terminus (DECHAT *et al.* 2008; WEBSTER *et al.* 2009). EXC-2, IFA-4, and IFB-1 do not have a CAAX domain, so do not appear to be bound to the apical canal membrane through the same mechanism as lamin. Other intermediate filament proteins such as keratin and vimentin form such meshworks linked to mammalian cell cytoplasm through a scaffolding protein, Plastin1 (IWATSUKI *et al.* 2002; GRIMM-GUNTER *et al.* 2009). Though *C. elegans* does not have an obvious plastin homologue, it will be interesting to see if a protein yet to be identified serves a similar purpose to link EXC-2 to the canal apical surface. Since EXC-2 retains its location in the canal and meshwork appearance when *ifa-4* and *ifb-1* are mutated and knocked down, respectively (Fig. 10), EXC-2 appears to be located to the apical membrane independently of these other two IF proteins. All three proteins are necessary to prevent cyst formation, and overexpression of *ifa-4* does not rescue *exc-2* mutation; these results suggest that the three filament proteins provide complementary functions to regulate tubule diameter and length.

We present a working model of these three proteins at the surface of the excretory canal in Fig. 11. The three intermediate-filament proteins EXC-2, IFA-4, and IFB-1 are collocated at the apical surface of the canal. We do not know if they are linked as obligate heterodimers; as EXC-2 binds to IFA-4 in our pulldown assay, this heterodimer presumably makes up some of the filaments. But EXC-2 is properly placed even without IFA-4 function, so EXC-2 dimers, either homodimers of one isoform or heterodimers of two isoforms, are likely part of the makeup of the filaments in wild-type animals.

**Figure 11.**
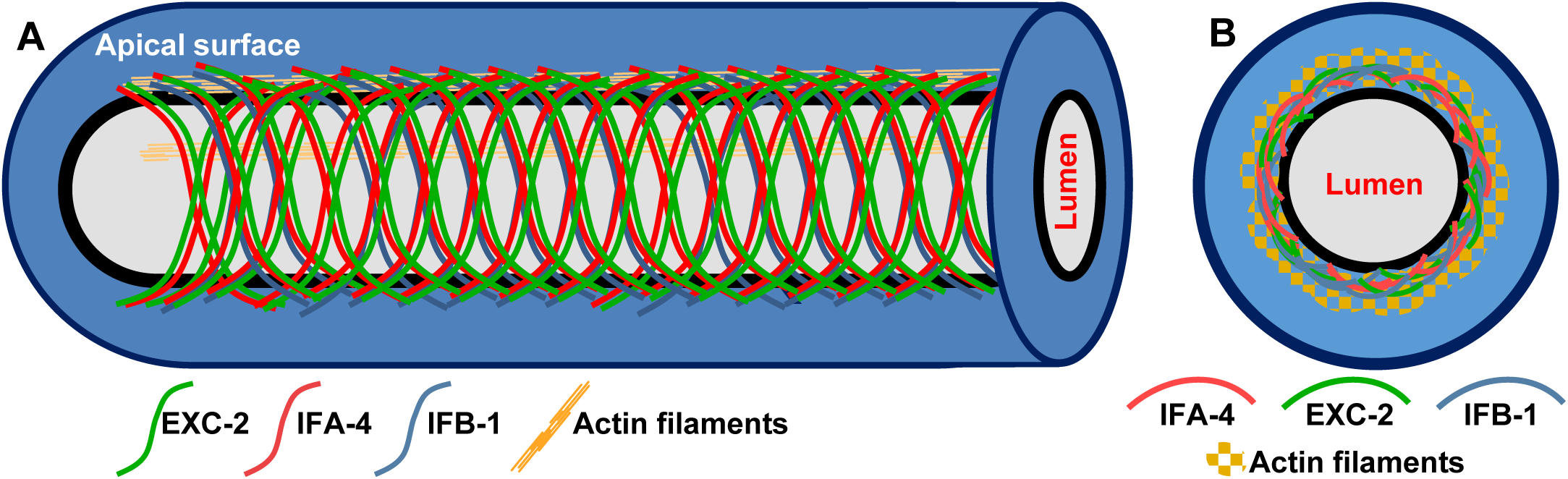
Proposed model of EXC-2, IFA-4, and IFB-1 in excretory canals. (A) Lateral section of the excretory canal, where intermediate filaments (green, red, blue) surround the apical membrane (black). Actin filaments make up the thick terminal web extending from the apical membrane and extending into the cytoplasm of the canals. Actin filaments are cut away in bottom half of drawing to show intermediate filaments more clearly. (B) Cross-section of the canal shows intermediate filaments forming a meshwork that surrounds the lumen surrounded by actin filaments.

The actin cytoskeleton forms a thick terminal web around the canal lumen visible in electron micrographs, and much thicker than the narrow band of intermediate filaments visible in the fluorescently tagged confocal micrographs here. The terminal web is tethered to the luminal membrane through the action of the SMA-1 β_H_spectrin (BUECHNER *et al.* 1999; FUJITA *et al.* 2003; PRAITIS *et al.* 2005)and the ezrin-radixin-moesin homologue ERM-1 (KHAN *et al.* 2013); we do not know if actin is closer to the membrane than are the intermediate filament proteins, but actin certainly lies outside of the layer intermediate filaments. Future studies on the *in vivo* interactions between these intermediate filament isoforms and proteins, and between these intermediate filaments and the actin cytoskeleton, should provide further insights into the ability of this network of proteins to provide the rigidity to maintain a firm luminal diameter while maintaining the flexibility to allow these narrow tubes to lengthen and bend during growth and movement.

## ACKNOWLEDGMENTS

We gratefully acknowledge the assistance of the KU Molecular Biosciences Sequencing Center. Other helpful discussions were provided by Stuart MacDonald, Meera Sundaram, members of the KU Genetics of Development Seminar, and members of the Baltimore Worm Club. We thank Verena Göbel for the fluorescently labelled *erm-1* strain. Some strains were provided by the CGC, which is funded by the NIH Office of Research Infrastructure Programs (P40 OD010440). Fosmid WRM0630A_E08 was the gift of the Max Planck Institute, Dresden, Germany. H.I.A. was supported in part by KU Graduate Research Funds #2301847 and #2144091. The Center for *C. elegans* Anatomy is supported by National Institutes of Health grant #OD010943 to D.H.H. E.A.L. was supported by National Institutes of Health grants #NS0090945, NS0095682, NS0076063, and GM103638.

